# Connections in the Dark: Social-Ecological Networks as a Promising Tool for Bat Conservation and One Health

**DOI:** 10.1101/2025.07.22.666214

**Authors:** Renata L. Muylaert, Cristina A. Kita, Michael Kriegl, Ahmad Bilal, Adeel Kazam, Chiara Scaramella, Emmanuelle Roth, Jon Flanders, Malik Oedin, Parfait Palamanga Thiombiano, Rida Ahmad, Sangay Tshering, Susan M. Tsang, Tigga Kingston, Tanja M. Straka, Marco A. R. Mello

## Abstract

Bats provide vital ecosystem services, including pest suppression and crop pollination. However, the increasing proximity of bats and humans is a growing conservation and public health concern with negative consequences for both sides. Mitigating these consequences requires integrative approaches like network science and the emerging field of social-ecological networks (SENs), which offer powerful tools to map and analyse complex social and ecological dynamics. Here, we synthesize how network approaches have been applied to bat research and conservation. Specifically, we: (i) assess the use of both ecological and social network analyses to study bats; (ii) identify network tools well-suited for SEN-based bat research; (iii) present a case study illustrating how SEN applications in bat research can inform conservation and One Health efforts; and, lastly, (iv) discuss key challenges and opportunities in using SENs to investigate the human-bat interface. Our review unveils a rise in network-based bat research from 2006 to 2020, followed by a post-pandemic decline. Nevertheless, across the 127 studies mapped by our review, only one applied an SEN lens. Finally, we suggest how applying some underexplored SEN tools to bat research could lead to novel perspectives, aiming to promote integrated strategies for the coexistence of bats and humans.

## Introduction

Bats are the second largest mammal order with nearly 1,500 species (*1*). Through their lives, bats interact with myriad other organisms, as more than 16,000 interactions have been documented in the peer-reviewed literature worldwide (*2*, *3*). Many of these interactions are vital for keeping ecosystems intact, being directly beneficial to people and their economies. For instance, bats are important pollinators for over 500 plant species, including economically important crops such as bananas, durian, and agave (*4*). They are also seed dispersers in tropical and subtropical forests and play a pivotal role as agents of pest suppression in agricultural and forestry systems across the world. Many species roost in large colonies resulting in accumulations of guano that can both fuel ecosystems, for example caves, and provide a source of fertilizer. Adaptations associated with their long life spans, flight, echolocation, hibernation, and immunology, among others, are sources of bio-inspired technologies and medical research (*5*, *6*).

Yet, some interactions with bats can also be negative. For example, bat colonies may be persecuted as a result of damage to commercial and backyard fruit crops (*7*, *8*), or because they are seen as the reservoir of viruses that can be pathogenic to livestock and people, such as rabies, Marburg, Nipah, Hendra, and, more recently, coronaviruses.(*9*) This duality places bats at the crossroads between conservation, economics, and public health, especially from the perspective of the UN’s Sustainable Development Goals (*10*) (such as life on land, and good health and wellbeing) and One Health(*11*) (a holistic collaborative approach connecting human health, animal health, and environmental health). In addition, bats epitomize “connections in the dark” – literally and metaphorically – since the ecosystem services delivered by those predominantly nocturnal mammals often remain unseen by the public (*12*). This invisibility fuels misinformation and heightens fear, further exacerbating conflicts and undermining solutions for global challenges, such as emerging diseases (*13*). Hence, relationships between bats and humans are dual, ranging from positive to negative (*14*).

Bats occur in different landscapes, from natural to human-modified, where in addition to their interactions with other taxa, these dual interactions can be found with a range of human stakeholders, being part of a variety of social-ecological systems. From rural to peri-urban and urban residents, farmers, orchard owners, decision makers to public health officials, different human actors, directly or indirectly, play a role in shaping perceptions about and behaviors towards bat communities and the ecosystem services they provide. Hence, to better understand the various complex interactions between humans and bats, state-of-the art and holistic approaches are needed to untangle the visible, but also hidden “connections in the dark” that shape the human-bat interface and their outcomes to promote ecosystem functioning including One Health (*15*).

There have been challenges to tease apart and quantify these connections, mainly because conservation and zoonotic disease mitigation efforts related to bats have traditionally focused on ecological and social concerns separately, often in antagonistic ways, with ecological queries focused on species, populations, or habitats (*16*), but not interactions (*17*). Although these approaches have yielded valuable insights and concrete results, they have often overlooked the interconnected webs, beyond the binary, in which bats are embedded (*18*). Those webs are woven by social and ecological relationships, involving human players who have the agency to deeply influence interactions, which can have positive or negative outcomes for both bats and people (*19*).

The study of how networks and their components are connected, or network science, can be potentially harnessed as a holistic approach to assess those webs, as it provides a conceptual and analytical framework to disentangle complex systems (*20*). Within the study of social-ecological systems, the framework of social-ecological networks (hereafter, SENs) offers a valuable approach using network science for exploring interconnections across the social and ecological domains (*21*). SENs map and describe the relationships both within and between the ecological (e.g., a food web) and human social domains (e.g., a governance network) using network metrics and analyses. For instance, SENs can be modeled from ecological networks focusing on animals, plants, fungi, or microbes, and their interactions with different players in human societies(*22*), or from interactions between the ecological and social domains of different environmental challenges (*23*). This transdisciplinary approach is highly valuable, because ecological networks are frequently reconfigured by human actions, which themselves are structured by social interactions across levels of societal organization and governance. Conversely, human societies may themselves be structured by the distribution of natural resources available to them. SENs, therefore, aim to bring together those multiple dimensions to help investigate the interplay between social and ecological processes (*24*). Over the last 20 years, SENs have been predominantly used in common pool resource management and to understand the complex relationships between social systems and ecological processes (*25*, *26*). For instance, researchers found that aligning social and ecological networks improves wildfire management, especially when partnerships reflect shared risks (*27*) or that social-social links are less strong in urban than rural areas when it comes to urban water and greenspace management (*26*).

Network science, with deep roots in mathematics, has evolved into a dynamic interdisciplinary field with applications in a wide variety of disciplines, including ecology and social sciences (*28*, *29*). Its rise in bat research highlights the importance of understanding the intricate connections between ecological and social factors in zoonotic disease transmission and ecosystem services (*21*, *30–32*). However, despite growing interest in this approach among bat researchers, its full potential remains untapped. Narrowing down to the subfield of SENs, there are exciting avenues to be explored. Applied at the human-bat interface, SENs could help us shed light on the hidden connections between bat ecology, ecosystem services, conservation, economies, and public health (*24*). This framework would help connect the dots between data and ideas, qualitatively and quantitatively, to develop more adaptive, context-specific, and comprehensive strategies to secure sustainable bat populations and healthy human economies and populations (*33*),(*^34^*).

Here, we discuss a promising pathway to achieve this goal through the identification and application of tools from network science suitable for studying the human-bat interface, by means of a systematic review and bibliometric analysis divided into four steps: (1) We begin with a research weaving approach (*35*) aimed at synthesizing knowledge about network science applied to bat research, focusing on gaps in social and ecological networks (SEN). (2) We then review and highlight promising concepts and tools from network science that remain to be explored in more depth and could help us reach a new level of insight into SEN research applied to bats; (3) Then, considering the results obtained in the two previous steps, we present a case study with varying contexts of human-bat interactions to illustrate the potential of SENs for bat research, conservation, and One Health. (4) Finally, we discuss challenges and opportunities in using SENs to investigate intricate relationships at the bat-human interface.

## Results

### How have social and ecological network analyses been used separately and jointly to study bats?

From our review of the literature pertaining to bat studies that used a network approach, we found that the earliest publication on the topic was from 2006 (*36*). Following the first review process, a total of 831 studies were discovered and narrowed down to 29, all of which consisted of non-SEN studies up to 2016. The updated period search generated 322 articles from 2016 to September 2024 that included screening of titles and abstracts. After applying our first stage inclusion criteria (studies that used analytical methods directly derived from network science including bats), 126 articles remained in our data set and were included in the second stage. Finally, when the methods and results sections were read, only the articles that met our second inclusion criteria were included in our review (studies that reported analytical methods directly derived from network science). This resulted in 100 papers being considered suitable for our review (Figure S3). Out of these 100 articles, two were also retrieved in the previous review, Webber et al. (*37*) and Zarazúa-Carbajal (*38*). Therefore, the combined counts from both searches (i.e., 100 and 29, minus two repeated studies) resulted in 127 studies linked to bats and network science. Interestingly, only one study, Kading and Kingston (*39*) applied a network approach using a bibliometric review that focused largely on social aspects of interdisciplinary collaborations (authors as social nodes and collaborations as links) related to bat zoonotic risk research. Because of its connection between a social network and ecological research outcome, we view this study as modeling a type of SEN. ‘Full SEN’ studies (SEN Type III) (*24*) were not captured in our review. From the 127 studies, we found an increase in the number of publications from 2016-2021, but a slight decline after 2022 (Figure 1A).

**Figure 1.**
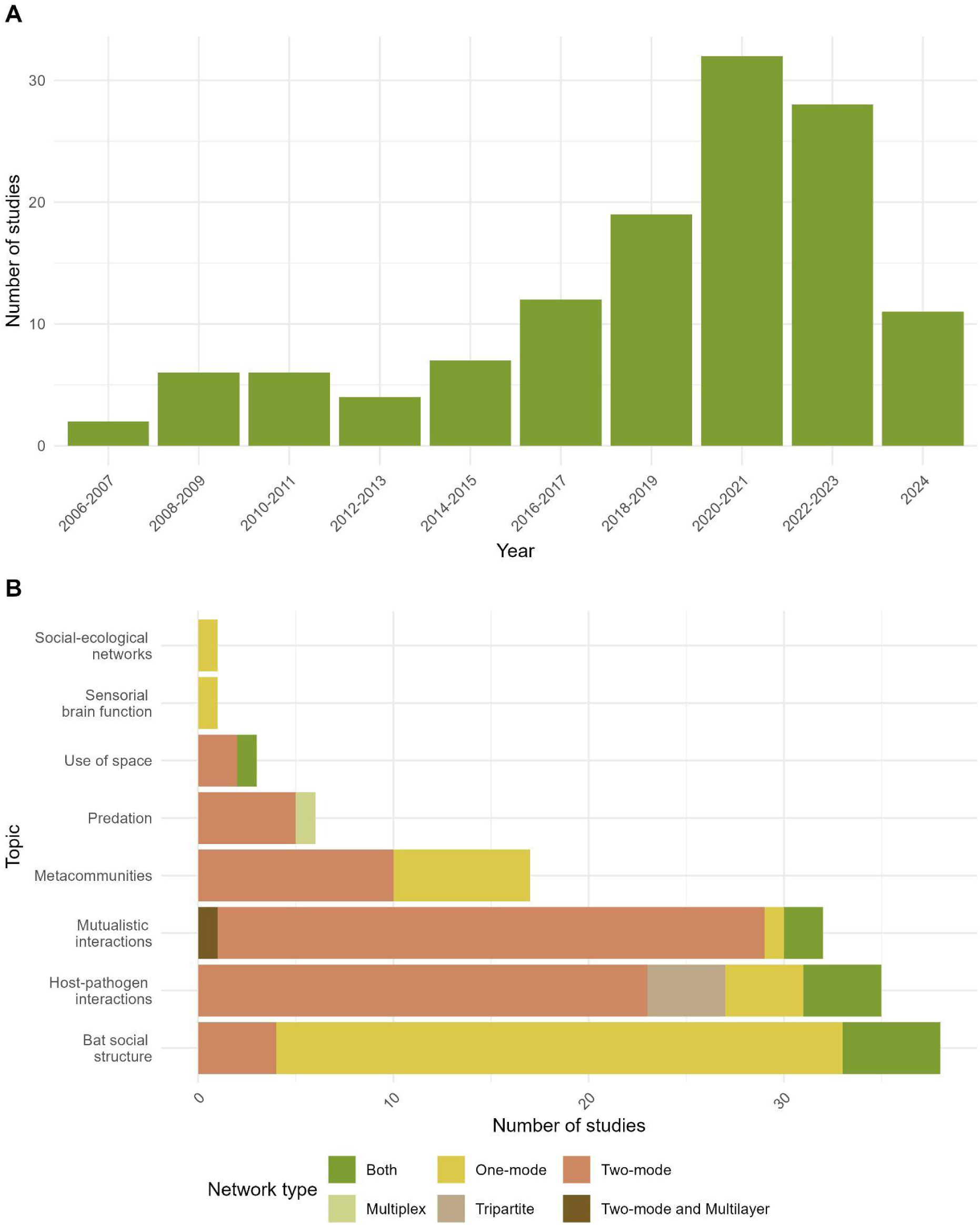
A. Temporal growth in bat studies that applied a network science approach up to the year 2024 (up to September 24th). B. The number of bat studies with a network approach per research topic. Bar colours indicate the type of network structure investigated. See all definitions for network types in Table S2.

We classified the selected publications according to main topics. Social interactions, mutualistic interactions, and host-pathogen interactions were the most frequently studied, (Figure 1B). Regarding interactions, the biological meaning of nodes and links varied considerably (Figure 1B, Table S4). For example, in studies on ecological systems at the community level, nodes were bat species, whereas for studies on bat populations, nodes represented individual bats. Depending on the topic being investigated, links in the studies represented interactions of frugivory or social relationships between bats, for instance.

The studies considered in our review varied in terms of topic, network type, and data type, with the most common topics investigating social structures among bats, thereafter referred to as “bat social structure”, followed by host-pathogen interactions and mutualistic interactions (Figure 1B). The least investigated topic was sensorial brain function related to social cognition in echolocating bats, modeled as functional brain networks. Most network types assessed were weighted (N=82), followed by binary (N=44) or both (N=7) (see glossary on Table S2 for Network Science terms). Despite their great potential, our review showed that SENs receive little attention in bat research, with only one study of this type in our review.

Collaboration was intense in bat studies with a network approach, with 541 individual coauthorship links and varying degrees of strength and repetition (average degree/collaboration number = 8, mode degree = 7, Figure S4). Individual co-authorships varied from 1 to 53 (Figure S5). The Neotropics and the Global South led the way in terms of the number of studies published (N=90). Several countries hosted only one study in addition to multi-country investigations throughout Europe and Central America. For single country and named multi-country investigations, there were multiple geographical research gaps, especially in Africa, the Middle East, Eastern Europe, and Southeast Asia (Figure 2). Brazil and the United States led the single country studies (N=20 and N=11, respectively), followed by Colombia (N=8) and Mexico (N=8). Multi-country studies investigated more than one country but had no global coverage (e. g., data from four countries, Neotropics, or North Africa, Figure S6). The study marked as *in silico* investigated vampire bat using simulated data (N=1) and was developed at Tohoku University, Japan (*40*).

**Figure 2.**
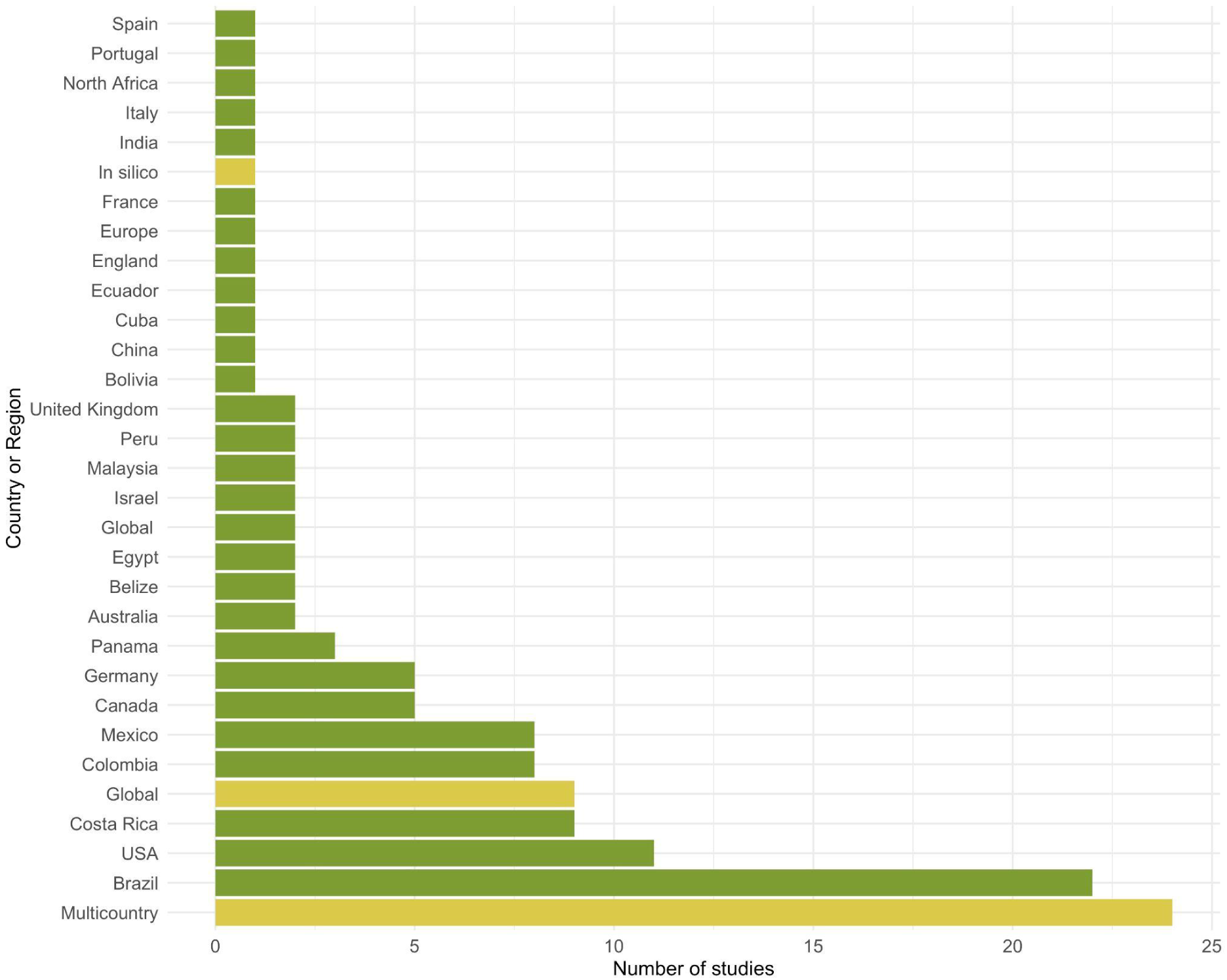
Number of ecological studies on bats with a network approach per geographic scope. Green bars represent data for a specific country, while yellow bars indicate data not associated with any particular country or multi-country efforts.

### Promising tools from network science to boost the SEN approach in bat studies

After a thorough review of the literature (see review metadata on Table S1), we identified promising tools from network science that remain under-explored, considering the tools identified in the reviewed papers and our experience with network science. The range of network metrics and tools used in the reviewed papers are summarised in Table S2. The top five metrics explored were nestedness (11.4% of studies), modularity (10.9% of studies), degree (9.4% of studies), specialization (4.7%), and connectance (4.1%). Sixty metrics were only mentioned once, including assemblage structure, a variety of connectivity metrics, and multilayer metrics.

As our research weaving approach shows, most studies so far have relied on networks evaluated as unipartite or bipartite (see glossary in Table S3), leaving indices derived from multilayer networks out. The first path we could take is delving deeper into assessments of **network structure** (also known as **<u>topology</u>**), mainly by expanding the types of metrics used to operationalize ecological and social concepts. In addition, while traditional topological metrics provide an overview of system-wide properties, they often miss finer details, which could be readily used as proxies for key social-ecological concepts (*24*, *33*, *41*).

Conventional topological analyses focused on bats often investigate trivial **<u>emergent</u> <u>properties</u>**, such as **connectance, nestedness, and modularity** (*34*). Trivial, in technical terms, means that they are based only on the definition of each organizational level (i.e., the entire network, its layers, its modules, its **motifs**, or its dyads) (*42*). For instance, nestedness is a trivial emergent property that is defined only for the entire network and not to its nodes separately. A more ambitious goal that has only been tangentially aimed for in social and ecological network studies is to delve into strong emergent properties that stem from complex, nonlinear interactions that cannot be directly inferred from the system’s components or definitions (*43*). This phenomenon is called **emergence**, and the key to studying it in the context of SENs is **criticality** (*44*, *45*). Criticality refers to qualitative changes occurring along system development, which have been used to assess when a network truly becomes complex (*46–48*), reaching network-level analysis and beyond the analysis of substructures of SENs, such as **<u>motif</u>** patterns (*49*). A classic example of strong emergent properties are characteristics such as hardness and conductivity, which tell apart carbon allotropes like graphite and diamond. In SENs, by examining how environmental conditions and social actors influence emergence in networks at the human-bat interface, researchers could uncover when critical ecosystem services, such as crop pollination, emerge from complex interactions and how they can be maintained or restored (*31*, *50*, *51*) or when critical thresholds of contact rates may disproportionately increase pathogen spillover risk.

Considering that through emergence different archetypical topologies arise in networks observed in natural, rural, and urban environments, it is also important to assess topology within a coherent theoretical framework. The study of SENs could benefit by harnessing a novel framework, the **Integrative Hypothesis of Specialization** (IHS) (*52*), which might also help identify key mechanisms and factors that regulate network assembly. IHS can help explain how heterogeneity in resource use drives a compound topology (modular with nested modules) in ecological networks (*53*). In doing so, IHS could help better understand which mechanisms shape critical SENs of interest. For instance, it could help detect the threshold at which crop pollination by bats breaks down in a given region, considering the resources available to bats. Because SENs can identify influential social actors within a system, such as key individuals or knowledge brokers, they can also offer valuable insights into stakeholder dynamics relevant to policy development (*54*).

Working within the realm of topology, another frontier to consider in SENs is **multilayer networks** (*55*). By definition, a SEN has at least two layers: one ecological and another social. Nevertheless, SENs can be modelled in much more complex ways, considering different ecological and social relationships that bind together the nodes representing humans, bats, and other organisms. Multilayer networks are a relatively new subdiscipline within network science (*56*), which has been growing quickly, drawing the attention of social scientists and ecologists (*57*). By exploring multilayer concepts and their proxies, bat researchers could, for instance, investigate different kinds of cohesive subgroups (*58–60*), like guilds and functional groups (*61*), that are composed not only of bats, livestock, or wildlife, but also of humans and their activities. By expanding the multilayer approach to SENs, we could also gain insight into how social links affect ecological links and vice versa. For example, if a novel zoonotic outbreak, associated with bats, affects ecosystem services, the potential result could hinder food production in a rural community. Fortunately, the number of multilayer tools available is rapidly increasing, and a myriad of metrics are ready to be explored (*62*).

There are also ways to analyze the relative importance of a node to the structure and dynamics of one or more layers in multi-layer networks (also known as centrality) (*63*). Multi-layer centrality applied to SENs could, among other possibilities, help expand and operationalize the concept of **keystone** species or components (*64*), needed for the study of complex ecological, economical, and sanitary relationships at the human-bat interface, also based on the concept of human’s as **hyperkeystones** (*65*). In bat studies, these analyses can help identify species that play critical roles in delivering ecosystem services (*66*) or acting as reservoirs for zoonotic pathogens (*67–69*). For instance, a bat species with high betweenness centrality might be critical for linking pollination and pest suppression (*70*) or promoting pathogen spillover (*71*). Similarly, a social (human) actor with high **betweenness centrality** might be a key actor when it comes to bat conservation. Conservation and restoration efforts targeted at hubs and connectors could optimize outcomes (*72–74*).

### Exploring SEN applications to bat research in conservation and One Health

For facilitating the uptake of SEN by bat researchers, we propose a stepwise guideline to study human-bat interfaces, with a case study as example (Table 1). Figure 3 provides a glimpse of the envisioned system that can be evaluated under a SEN perspective, with tangible implications for bat conservation and public health. We illustrate a published work with different contexts of human-bat interactions and reimagine its investigated system as case study to demonstrate how social and biological entities at the human-bat interface could be assessed together following an SEN approach. The case study explored the potential for using bats as medicines across different regions in the world, based on secondary data (Figure 3), where regional to global patterns can be explored as SENs. In Tackett et al. (*75*), a bat-medicinal use system can be explored as networks with metrics and models to be applied to all in-network, multi network and external-to-network components that could benefit One Health and bat conservation (Figure 3, Table 1). A few unexplored metrics for this system are related to operationalising the identification of key actors that could reduce the harm that animal trafficking causes bat conservation (for instance, connectors in the network and hubs), and local healers that could be informed about potential zoonotic risks associated with bat handling and use (through exploring betweenness centrality variation in the different social actors who are connected to ecological nodes).

**Figure 3.**
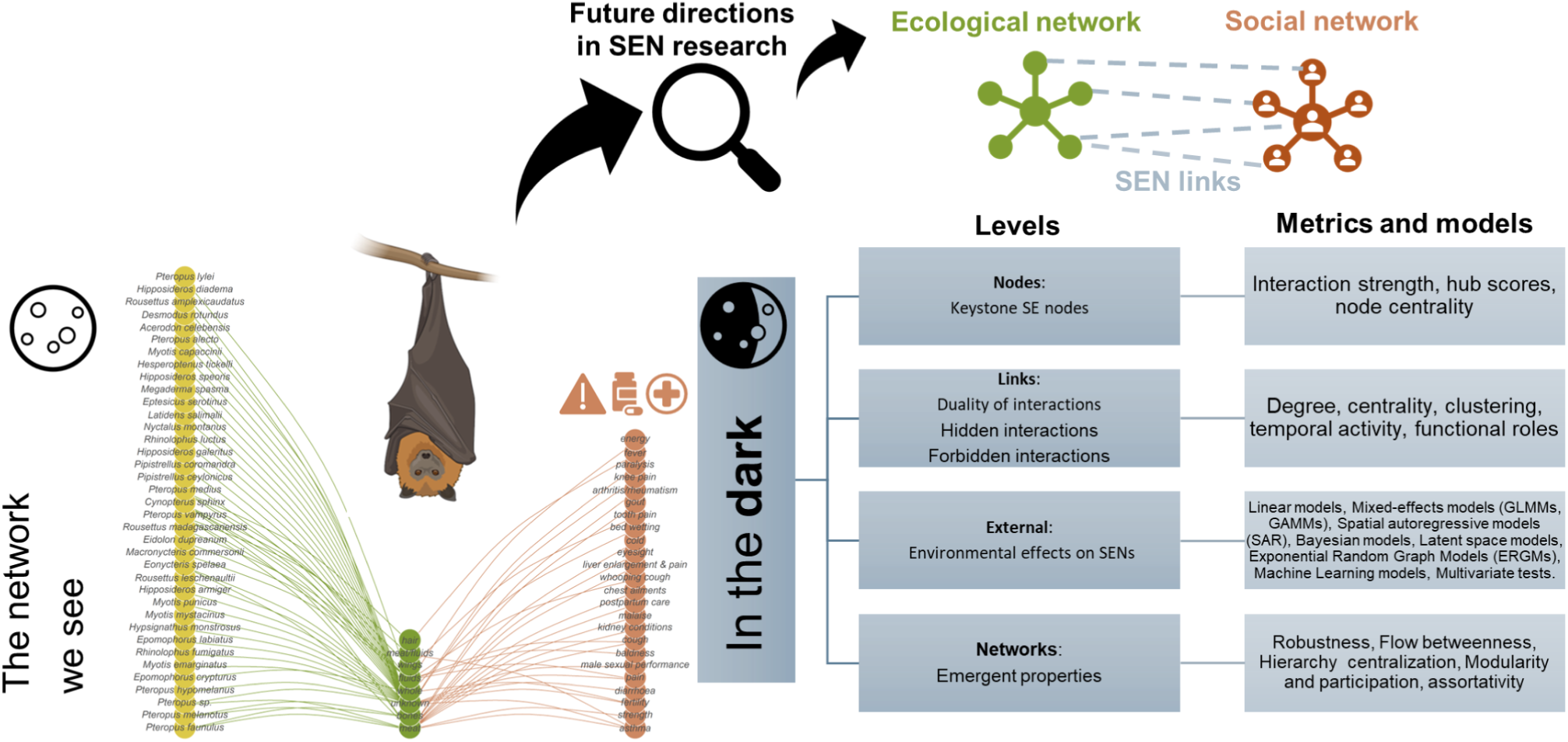
An example of secondary data to be yet explored in future SENs-bat research. The network displays medicinal uses of bat species (nodes on the left) by humans across 83 countries and territories as a social-ecological network (SEN). The network depicts different types of nodes as bat species (left), bat body parts harvested (centre), and illnesses/conditions treated directly linked to medicinal use by people (nodes on the right). The figure was composed with a bat silhouette from BioRender. Data source: Tackett et al. (*75*).

**Table 1.** A general step-wise guide to construct an SEN project related to bat conservation and One Health, envisioning a study case. Methods for this framework are expanded from Fig. 3 and (*77*).

| Step | Description | Study case: Envisioning a project based on the system investigated by (Tackett et al. 2022) |
| --- | --- | --- |
| 1 | Define a clear research question for social-ecological interdependence related to bats and define the limits of your network (bat assemblage, protected area, global?) | List target places where bats are used, cultures, flow of trade, official lists. Check for ethics at every step due to sensitive data prior to project commencement. |
| 2 | Identify ecological (e.g., species, habitats) and social (e.g., individuals, organizations) nodes relevant to the research question and your study system. Evaluate the need to aggregate nodes from individuals into groups. Group the variables into ecological and social. | Group variables, taking into account that human actors must be deidentified. Bat species may also have regional variation as subspecies and varying levels of extinction threat. Data from large global databases can be used to assess species status and distribution. |
| 3 | Think about ethics all the way and apply for associated research permits. Identify unidirectional linkages and the effect variables (outcome variables). Refine and tidy up your envisioned format for adjacency matrix(es) and dependencies. | Establish pathways between social and ecological links, investigating trade chains, sources and market outlets. Check how duality can play a role in different communities as bats are viewed positively in a community and still be exploited. Think about quantifying these variables on an edge list. E.g.: Assigning values for positive conservation actions in place and negative values for antagonistic social-ecological interactions such as illegal trade. |
| 4 | Organise your plan for data collection of interactions (77) for operational steps for data entry. including how you will attribute values to your SS, SE, EE links. | Ultimate outcome variable or effect: detangle dual use of bats, dual perceptions of bats, potential ecosystem services and disservices (if applicable). |
| 5 | Data collection: Quantify/Encode/Score all the data nodes based on research questions, potential your hypothesis and premises. | After ethics approval, collect the data. Consider potential lack of coverage for sensitive nodes. |
|  |  | Expect that discretisations of data might happen for certain nodes or even binary interactions (presence and absence). |
| 6 | Data wrangling: organise, tidy up your network data and metadata for analysis. | Identify pathways to be investigated and complex/hidden/latent interactions to be expected, for instance hierarchical dependencies on institutional constraints (could be measured through centralization metrics), motifs and connectivity patterns (e.g. economic demand, lack for law enforcement in peripheral nodes, and motifs formed by shared cultural norms). |
| 7 | Analysis: select the level of complexity and kind of network analysis (node level, link level, network level). Analyze your network using relevant network metrics for your research question. | Work on visualization and module detection. Attempt to understand criticality through distance metrics. Connectivity analysis can inform about central functional roles. |
| 8 | Research questions answered and communicated. After communicating your findings, can you update the network with new data / learnings and discuss future relevant contexts (depending on project duration and funding)? Observe what is confirmed or predicted by the network reasonably and what further insights / complexities / hidden causality emerge from the network (i.e. Emergence). Discuss critical node role and system change during transient dynamics (i.e. Criticality). | Make the network available to relevant stakeholders, think about project continuity as data availability evolves. Consider adding a temporal dimension to the analysis for future projects. Discuss connections in the dark detangled, forbidden links and emergent properties of the system. What nodes/actors or link weights/directions would need to be changed to improve One Health and bat conservation? |

In terms of diving deeper in the social components of an SEN, having our study case as a lighthouse, changes in the dynamics of cultural views between elders and youth may contribute to reduced consumption of bats. In that sense, information could be viewed as links in the SEN investigated. It is important to apply social-ecological approaches so conservation measures do not drive the loss of Traditional Ecological Knowledge (TEK) and Indigenous Knowledge (IK), and contribute to overall social-ecological resilience, which is the capacity of interconnected social and ecological systems to deal with external stressors and disturbances without losing their core identity and functions, ensuring the continued provision of essential benefits that a community obtains from the ecosystem (*76*). The importance of bats to social-ecological systems is far from fully understood by researchers. Also, the potential for transmission and pandemic potential of known bat-borne pathogens may not be fully known by different cultures and populations, and potential for risk is generally unknown to researchers as well. SENs can be used to explore relevant links between biodiversity dimensions and health, and ideally, indicators for Conservation and One Health through SENs could be proposed. Conservation information provides opportunities to communicate on the ecological importance of bats while noting the potential risks of consuming them. SENs can support opportunities in community development, education, ecotourism, economic integration, values and beliefs, and integrated land-use planning.

## Discussion

### How have social and ecological network analyses been used separately and jointly to study bats?

Our review highlights a general growth in bat studies using a network approach, but a specific gap related to social-ecological networks. There was a notable surge but then a decline in publications following the onset of the COVID-19 pandemic, which could indicate a shift of focus or funding restrictions. The Neotropics emerged as a leading region for contributions, reflecting a significant focus on bat research and network science in this biodiverse region. Collaboration was a defining feature of this body of work, with over 500 recorded author partnerships heavily focused on multi-country study regions and the Global South, varying in intensity and recurrence, underscoring the importance of interdisciplinary and international efforts.

First, we wish to discuss the drop in publications observed during the COVID-19 pandemic. With 2020 being the most disruptive and uncertain year of the pandemic and 2021 being the deadliest, 2022 still recorded most cases (many being Omicron variant) (*78*) but with declining mortality and the start of the onset for a new post-pandemic ‘normal’, which might reflect this late pattern in studies declining. Importantly, pandemic years notably increased publication rates in bat-pathogen research (*79*) since there was concern about bats and disease ecology, and researchers working through existing datasets and unfinished projects since lockdowns interfered with fieldwork. We also aggregated data in 2-year bins, so this pandemic disruption in the trend needs to be carefully evaluated as our last period does not end in a full 24 months bin.

Anyway, despite the general growth – and pandemic fall – in bat studies with a network approach, SENs remain widely underutilized. We outline the benefits that network approaches, particularly social-ecological networks, can provide to bat research, such as revealing interactions and feedback among species, environments, and people. A network approach also helps identify leverage points, e.g., important (social) actors in a network, manage trade-offs, and design integrated, cross-sectoral solutions that are crucial for effective bat conservation and One Health. We also argue that particularly SENs may hold significant potential for addressing complex challenges such as within the One Health framework and for conservation, and so this specific approach should be further explored in future studies.

Previous research about the relationship between humans and bats has mostly concentrated on conflicts between humans and bats and concepts such as attitudes, knowledge, or beliefs (*14*). These studies typically explore how much people like or know about bats, whether they inspire joy or provoke fear and disgust, and how these thoughts or feelings translate into behaviors towards bats. From a network perspective, this is a form of social-ecological link, such as how much joy or fear bats engender in people (with bats as ecological nodes, humans as social nodes, and the emotions as links between them). Building SENs that capture more direct impacts of human actions on bats might formalize links between them based on habitat management actions, hunting, persecution, or other major drivers of bat population declines (*80*). However, what is largely missing in this research is understanding how, for instance, social actors influence each other and how this translates into behaviour towards bats. This is where a SEN approach comes in.

Despite the growing interest in human-bat relationships (*14*), our review shows that there was only one study that adopted a SEN approach, as defined by our search criteria. The sole study in our review that we consider has a social-ecological approach (*39*), resulting from a literature review, and having a comprehensive network perspective about coauthor relationships in bat disease research. Its authors (*39*) identified via a coauthorship network only a few interdisciplinary collaborations between disease and ecological researchers, but critically those that existed were underpinned by a common goal.

In this spirit, SENs have the potential to similarly unite and integrate perspectives from different disciplines, particularly those seeking sustainable outcomes for both wildlife and human populations. This is one of the strengths of SENs, understanding where more resources are needed (*81*) or should be allocated. In the aforementioned study, also showing an imbalance in interdisciplinary collaborations. Referring to the definitions of Kluger et al. (*22*), this would be considered a type II network because only social networks were included, but what brings the social actors together is the study of bats. Interestingly, 2006, when our review found the first bat studies with a network approach, is also the year when one of the first publications more or less related to SENs appeared. This starting point reveals a clear lag in adopting SENs in bat studies. This may be because full SEN studies require extensive relationship building and collaboration between social and ecological experts. This is an ongoing challenge in bat conservation. Through our review, we tend to agree with the critique by Sayles et al. (*41*) that most SEN research lacks balance. While their review emphasizes the need for more involvement of ecological network experts in SEN studies, we propose that ecological network research on bats could significantly benefit from incorporating more social network expertise (we acknowledge that this perspective is primarily grounded in ecological research).

### Tools from network science that could be integrated into an SEN approach focused on bats

While network science is gaining traction in bat-related research, much of its analytical potential is yet to be explored. Expanding beyond basic topological metrics to include **emergence**, criticality, and multi-layer networks would enable researchers to answer complex questions about SEN dynamics. This deeper understanding is essential for developing effective strategies to address critical biodiversity conservation and public health challenges.

### Exploring SEN applications to bat research in conservation and One Health

SENs can be also used to understand the human-wildlife interfaces (Figure 3), public perceptions, transboundary diseases, and ecosystem services. If SEN analyses are to guide interventions, there are a number of considerations and caveats. For instance, addressing the use and perceptions of bats and their interactions with different cultures can be challenging to factor when exploring the applications of SENs or implementing conservation, particularly if bats are regarded as bush meat or used as a ceremonial food resource (*75*). As explored in our study case, the variety of bat species harvested for ceremonial purposes also poses challenges for conservation and makes it difficult to regulate. In light of this, beneficial outcomes can be explored where bats are regarded as sacred or important by indigenous cultures. Moreover, there is some evidence of win-win solutions when the social-ecological approach is used to identify win-win solutions to endure future anthropogenic and environmental challenges. An example of this is the mapping of successful conservation models including ample engagement with Indigenous Peoples and Local Communities of various cultures, also called socio-ecological ‘*hopespots’*, in the Amazon (*82*).

### Opportunities and challenges

Network science has the power to reveal the underlying structure of ecological and social systems at multiple scales. SENs have the power to advance our understanding of how bat conservation can be improved while also benefiting public health. There are further opportunities for bat conservation to use network science, for instance, to understand how interactions among decision makers impact bat populations. Based on our analysis, we identified knowledge gaps and biases, and point out paths for future studies based on a study case and a proposed SEN framework, identifying where efforts and resources can be better invested considering challenges and opportunities around the implications and trends on using social-ecological networks as a tool for bat conservation and public health.

Using network science and the SEN approach offers immense potential for addressing challenges in bat conservation, public health, and climate resilience. Key benefits include identifying critical ecosystem services provided by bats, understanding complex interactions between bats, humans, and livestock, and mitigating zoonotic disease risks. By enhancing collaboration across sectors (e.g., Quadripartite, One Health Monitoring, and co-design initiatives), the SEN approach promotes resilience in both human and environmental systems, fostering sustainable practices across watersheds, cities, and countries.

However, implementing the SEN approach comes with challenges. Methodological constraints, climate-related stressors, and the complexity of SENs – such as interpreting multiplex layers, noise, and hierarchical effects – can hinder progress. Additionally, prioritizing conservation efforts in areas where bats, livestock, and human interactions create complex nexus points remains a challenge. Yet, opportunities abound in leveraging SENs to mitigate climate change (e.g., bat-proofing roof initiatives in the Netherlands and elsewhere) and promote humane solutions for bats, fostering social-ecological resilience at the human-bat-livestock interface.

Cross-disciplinary collaboration is essential to overcoming these challenges, but barriers – such as global disparities, limited funding, and small research groups – complicate mobilizing efforts. By utilizing high-tech methods, citizen science, and open data, tangible collaborations and applications can emerge, enhancing resource allocation and funding for bat conservation and public health initiatives. Translating patterns, theories and mechanisms into actionable policies, lack of roadmaps that help bridge knowledge syntheses to actionable science and policy are pervasive (*83*, *84*). Contextualising work on areas where pervasive negative cultural beliefs around bats exist and/or where conservation is not a priority is paramount (*85*).

Moreover, public engagement is critical in shifting perceptions, especially in areas with negative cultural beliefs about bats. By focusing on community-based conservation, promoting positive perceptions, and leveraging media campaigns, the SEN approach can reveal and influence public behavior and support for biodiversity conservation (*86*), One Health (*11*) and Sustainable Development Goals (SDGs) (*87*). Ultimately, translating scientific patterns and theories into actionable policies in health and conservation will require overcoming knowledge gaps and ensuring that strategies are tailored to different cultural and regional contexts. Finally, communicating the interdependence of components in SENs can empower communities that have a closer connection with nature and bats (*88*) and point out directions where positive perceptions are prone for growth (*89*).

## Methods

Here we reviewed how network science has been used in ecological and social-ecological (there are often both names: socio-ecological and social-ecological; we mostly use the latter) networks that focus on bats as a key component. We provide high-level information on link types, study distribution through a systematic review and bibliometric mapping Nakagawa (2019) (*35*) of SENs and non SENs. First, we surveyed the literature on Scopus using the following search terms or tokens on a pre-screening phase: socio* OR socia* OR *ecolog* AND bat OR chiroptera AND network* OR graph\** (322 Scopus, a narrower range of papers), and a complementary comprehensive search using *bat OR Chiroptera AND network* OR graph\** (1,856 articles). Based on a pre-screening assessment of search results (Supplementary Text S1, Figure S1, Figure S2), we deemed the first search term as adequate for screening.

We restricted the searches in the Scopus database from 2016 to October 2024, expanding a previous review that focused mainly on ecological dimensions (*34*). Eligible articles resulting from the search were all original articles in English that could be retrieved online or per personal author request. Articles were screened by reading titles, abstracts, and key-words. Previous reviews on social-ecological networks have for instance focused on human-biodiversity interactions in general (*22*) or sustainability science (*41*), introducing the definition of various network types, such as Type I, II, and III (*22*), or non-articulated to fully articulated (see all definitions on Table S3) (*41*), depending on how ecological and social dimensions are connected in their network representation. In our literature review focusing on bats, the main criteria for inclusion during the screening step was the application of analytical methods directly derived from network science to some aspect of bat biology. We did not consider studies coming from other theoretical and analytical frameworks, such as multivariate statistics, niche overlap analysis, or fundamentally/purely descriptive interactions (like a bat feeding on human blood). The first stage narrowed the papers down to 124 papers through titles and abstract screening following the criteria that the study must be related to network science research including bats. The second stage focused on methods and results curated studies that reported analytical methods directly derived from network science. Eligible articles that could be retrieved were then downloaded and organised in folders to be read by authors.

Article metadata and derived author collaborations were generated through a custom workflow using the screened output the search imported and analyzed using *litsearchr* (*90*) and custom functions. Since bat-SEN studies are much rarer than ecological network studies, we also screened the literature cited by the bat-SEN study to increase the range of search. In order to report searches, screened, retrieved, eligible and excluded articles, a PRISMA chart (*91*) was generated (Figure S3).

We extracted data including the list of authors, year of publication, country or regions where the study data comes from, network type (SEN, non-SEN), and topics evaluated in terms of biological meaning of nodes and links. We also evaluated whether the studies include data from the Global South and contrast Global South-based studies with non-Global South and global studies. Reviewed article metadata were used to build a collaboration network. The number of individual author collaborations and summary statistics were calculated to reveal the main patterns in the collaboration network (S4 Fig). Modularity was calculated using the Louvain algorithm (*59*), that optimally assigns a membership for each author to a cluster. Importantly, as network science can present itself with numerous jargons that can confuse cross-disciplinary teams, we present a general glossary in Table S2 and its consultation is advised for the reader that has just started to have contact with network science and SENs. The glossary supports the reader to understand our results and their applications in their own future projects.

### Tools from network science

Based on reviewed articles, the metrics and network science tools used and our own experience, we present and discuss tools from network science that could be applied in projects proposing an SEN approach focused on bats analytically, be focusing quantitative analysis and variables used for investigating network-level and emergent properties.

### Exploring SEN applications to bat research in conservation and One Health

From describing useful tools, we propose a general stepwise method for applying the SEN framework to bat studies. For creating our steps, we use as references the SEN frameworks and network science workflows, such as Straka et al. (*92*) and Kita et al. (*77*). We proceed to explore SEN research tools and directions based on one network available from the literature: fruit bat benefits and costs networks and medicinal use of bats by humans. All analyses were made in R 4.5.0 (*93*) using the packages *rcrossref* (*94*) and *igraph* (*95*).

### Opportunities and challenges

From our results, we then proceed to discuss opportunities and challenges of applying the SEN approach to scientific research projects that could inform Conservation and One Health actions.

## Acknowledgments

We are deeply grateful to the authors of all primary studies included in our systematic review, whose empirical work made our synthesis possible. RLM is supported by an Australian Research Council Australian Laureate Fellowship (FL240100037) funded by the Australian Government. This work is a product of the SEN Working Group of the Global Union of Bat Diversity Networks (GBatNet). GBatNet activities are supported by the National Science Foundation AccelNet Award Numbers 2020595, 2020577, and 2020565. Any opinions, findings, conclusions, or recommendations expressed in this work are those of the author(s) and do not necessarily reflect the views of the National Science Foundation. TMS acknowledges the CURT network funded by the Deutsche Forschungsgemeinschaft (DFG; STR 1714/2-1) and Jonathan Jeschke for the insightful and inspiring discussions on social-ecological networks, which significantly shaped her thinking.

## Data accessibility statement

Does not apply.

## Code accessibility statement

Processed data and codes in support of this publication are publicly available on GitHub and via Zenodo (TBD):

1. https://github.com/renatamuy/bibmap
2. https://github.com/renatamuy/tackett

