## Supporting Information for "Connections in the Dark: Social-Ecological Networks as a Promising Tool for Bat Conservation and One Health"

### Supplementary Material

**Table S1.** Metadata used for synthesizing knowledge about network science applied to bat research.

| Column | Description | Value (type and example) |
|---|---|---|
| **Number** | Study number | Numeric: from 1 to inf |
| **Reference** | reference (short name for the paper) | String: Muylaert et al 2013 |
| **DOI** | DOI without https:// | String: 10.1515/mammalia-2012-0103 |
| **Year** | year published | Integer: 2013 |
| **Global South** | Does the paper include data from global south? Possible values are Yes or No. Values possible for global south = YES : Africa, Latin America and the Caribbean, Asia (excluding Israel, Japan, and South Korea), and Oceania (excluding Australia and New Zealand). Source: UN Trade and Development | String: Yes |
| **Country_or_Region** | Country name(s) separated by [and], Region (when Neotropical etc) or global. We detangle this later with code. | String: Brazil |
| **Family** | Family names should be separated by [and]. If too many families like more than 4, just mention multifamily | String: Phyllostomidae |
| **Networks** | Column created. Values possible: Not-SEN, SEN1, SEN2, SEN3, SEN4 sensu Kluger et al. 2020. It considers a network of bat researchers or bat workers as an aspirational SEN. | String: Not-SEN |
| **Links_EE** | Link type(s) ecological processes. Categories are context-dependent and may grow as more studies are curated. | String: Mutualistic interactions |
| **Links_SS** | The social link(s) in the study. | String or NA: NA |
| **Links_SE** | The social-ecological link(s) in the study. | String or NA: NA |
| **Links_detail** | Detailed free hand tag for links | Frugivory and Seed dispersal |
| **Nodes_EE** | The ecological meaning of the nodes in the study. | String: Individual Sturnira lilium bats and plants |
| **Nodes_SS** | The social meaning of the nodes in the study. | String or NA: NA |
| **Mode** | One-mode, two-mode, both. | String: Weighted |
| **Weights** | Weighted, binary or both. | String: Two-mode |
| **Network** | General type of network metric used. In case of network models, we refer to them just as network | String: Centrality |

| Metrics Used | models |
|---|---|

Table S2. Tools and metrics from network science and its usage in our reviewed articles. Graph analysis here refers to studies who represented their systems as a graph

| Network tools | Number of mentions | Percentage (%) |
|---|---|---|
| Nestedness | 39 | 11.4 |
| Modularity | 37 | 10.9 |
| Degree | 32 | 9.4 |
| Specialization | 16 | 4.7 |
| Connectance | 14 | 4.1 |
| Betweenness Centrality | 8 | 2.3 |
| Complementary specialization | 8 | 2.3 |
| Graph analysis | 8 | 2.3 |
| Network size | 8 | 2.3 |
| Network structure | 8 | 2.3 |
| Closeness Centrality | 7 | 2.1 |
| Eigenvector Centrality | 7 | 2.1 |
| Centrality | 6 | 1.8 |
| Co-occurrence | 6 | 1.8 |
| Interaction strength | 6 | 1.8 |
| Network density | 6 | 1.8 |
| Degree Centrality | 5 | 1.5 |
| Network models | 5 | 1.5 |
| Robustness | 5 | 1.5 |
| Assortativity | 4 | 1.2 |
| Connectivity | 4 | 1.2 |
| Dyadic associations | 4 | 1.2 |
| Weighted degree | 4 | 1.2 |

| | | |
|---|---|---|
| Association strength | 3 | 0.9 |
| Betweenness | 3 | 0.9 |
| Communities | 3 | 0.9 |
| Graph density | 3 | 0.9 |
| Mean degree | 3 | 0.9 |
| Participation coefficient | 3 | 0.9 |
| Accessibility | 2 | 0.6 |
| Average path length | 2 | 0.6 |
| Clustering coefficient | 2 | 0.6 |
| Diameter | 2 | 0.6 |
| Interaction specificity | 2 | 0.6 |
| Node strength | 2 | 0.6 |
| Number of compartments | 2 | 0.6 |
| Relative degree | 2 | 0.6 |
| Assemblage structure | 1 | 0.3 |
| Asymmetry | 1 | 0.3 |
| Average degree | 1 | 0.3 |
| Average degrees | 1 | 0.3 |
| Between-module connectivity | 1 | 0.3 |
| Betweenness Centrality | 1 | 0.3 |
| Bridging index | 1 | 0.3 |
| Closeness | 1 | 0.3 |
| Clustering | 1 | 0.3 |
| Clustering degree | 1 | 0.3 |
| Clusters | 1 | 0.3 |
| Compound topology emergence | 1 | 0.3 |
| Core-periphery classes | 1 | 0.3 |

| | | |
|---|---|---|
| Core-periphery network structure | 1 | 0.3 |
| Core–periphery network structure | 1 | 0.3 |
| Degree assortativity | 1 | 0.3 |
| Degree centralization | 1 | 0.3 |
| Degree distribution | 1 | 0.3 |
| Density | 1 | 0.3 |
| Functional complementarity | 1 | 0.3 |
| Functional role | 1 | 0.3 |
| Generality | 1 | 0.3 |
| Homophily | 1 | 0.3 |
| In-degree Centrality | 1 | 0.3 |
| Interaction Strength | 1 | 0.3 |
| Interaction signal | 1 | 0.3 |
| Linkage density | 1 | 0.3 |
| Links per species | 1 | 0.3 |
| Maximum path length | 1 | 0.3 |
| Mean Clustering coefficient | 1 | 0.3 |
| Mean node betweenness | 1 | 0.3 |
| Mean path length | 1 | 0.3 |
| Multilayer Centrality | 1 | 0.3 |
| Network Centrality | 1 | 0.3 |
| Network clusters | 1 | 0.3 |
| Network degree centralization | 1 | 0.3 |
| Network diameter along node loss | 1 | 0.3 |
| Network stability | 1 | 0.3 |
| Niche overlap | 1 | 0.3 |

| | | |
|---|---|---|
| Node betweenness | 1 | 0.3 |
| Node contribution to nestedness | 1 | 0.3 |
| Number of transitive triplets | 1 | 0.3 |
| Ontogenetic switch | 1 | 0.3 |
| Phyllogenetic signal in network | 1 | 0.3 |
| Phyllogenetic signal on network | 1 | 0.3 |
| Quantitative Modularity | 1 | 0.3 |
| Robustness to extinction | 1 | 0.3 |
| Roost degree | 1 | 0.3 |
| Second-degree Centrality | 1 | 0.3 |
| Size | 1 | 0.3 |
| Specificity | 1 | 0.3 |
| Strength Centrality | 1 | 0.3 |
| Strength and betweenness | 1 | 0.3 |
| Symmetry | 1 | 0.3 |
| Transitivity | 1 | 0.3 |
| Trophic node composition | 1 | 0.3 |
| Vulnerability | 1 | 0.3 |
| Weighted nestedness | 1 | 0.3 |
| Within-module connectivity | 1 | 0.3 |
| Within-module degree | 1 | 0.3 |

**Table S3.** A comprehensive glossary of network science and social-ecological networks (SENs). In some cases, the same concepts are represented by different terms in each literature. Terms explained circularly in this dictionary are written in *italics*. Adapted from concepts defined in Mello and Muylaert (2020), Kluger et al. (2020), Sayles et al. (2019), and Barnes et al. (2022) (*49*). Not every concept defined here is necessarily mentioned in the main text.

| Term | Definition and usage |
|---|---|
| SEN Network Type 1 | Considers one type of node (either from the social or from the ecological realm) and one type of link (either SS or EE interactions). Since only one realm (social or ecological) is represented, SE links are not incorporated. |
| SEN Network Type 2 | Integrates two types of nodes (from both the social and from the ecological realm) and two types of links (SS and SE, or EE and SE links). Although the interaction of one realm with certain actors from the respective other dimension is conceptualized, no further links between the nodes of that other realm are considered. |
| SEN Network Type 3 | comprises two types of nodes (from both the social and from the ecological realm) and three types of links (SS, EE and SE) between these actors. |
| Social actor | Individual, group, organization, or institution that participates in and influences social interactions, behaviors, or structures within a network. In a network, a social actor is treated as a node. |
| Partially articulated network | Type II SEN (see *SEN Network Type 2*). |
| Fully articulated network | Type III SEN (see *SEN Network Type 3*). |
| Adjacency matrix | A matrix that defines which *vertices* in a *graph* are connected to one another by an *edge*. |
| Average path length | A metric of *connectivity*. Considering all small *paths* between all pairs of *nodes* in a *network*, the average path length is the arithmetic mean of the length of those small paths. For example, “six degrees of separation” is a concept related to how many social relationships, on average, separate any two persons in the world. |

| | |
|---|---|
| Betweenness | A metric of *centrality*. The proportion of *small paths* in the *network* in which the *node* is present. For example, a bat species that feeds on flower species of different *modules* in a network is expected to have high betweenness, as it is a bridge between regions of the network. |
| Binary | A *link* whose value is either 0 or 1. A *network* based on presence or absence of links is called binary. For example, a bat species visiting or not visiting a plant species in a pollination network. |
| Bipartite | A *network* that is divided into two *sets* of *nodes*, in which *links* may exist only between nodes of different sets. For example, a network formed between nectarivorous bat species and the plant species visited by them: bats may visit plants, but not other bats. Synonym: *two-mode network*. |
| Centrality | The relative importance of a *node* to the *topology* of its *network*. There are many different concepts of centrality focused on different aspects of a node's importance. For example, the *degree* of a node is a kind of centrality. |
| Closeness | A metric of *centrality*. The average number of *small paths* that separates a given *node* to any other nodes of the same *network*. For example, a bat species that visits a set of flower species, which are visited by many other bats in a network, is expected to have high closeness. In other words, the bat species has a very common niche. |
| Combined | A *network* with two or more predominant *topologies*, usually hierarchical to one another. For instance, a network that has a *modular* topology, and whose modules that are internally *nested*. |
| Community | In network science, a *community* is a cohesive subgroup of *nodes* or *links* in the *network*, which are more strongly related to one another than to other nodes or links of the same network. Communities are also called *modules*. |
| Complementary specialization | A *connectivity* metric. It assesses how much the *nodes* in a *network* establish unique *links*. For example, in a bat-plant network, if each bat species visits a different *set* of plant species, and niche overlap is close to zero, then the network scores high complementary specialization; in other words, the bat species play complementary functional roles. |
| Complex system | A *system* whose properties cannot be inferred only from its *elements* or *relationships* is called a complex system. The properties of a complex system emerge from its assembly, and so they are called emergent properties. For example, a *network* may be *nested*, but a *node* may not. |

| | |
|---|---|
| Connectance | A metric of *connectivity*. The proportion of *links* observed in a *network* in relation to the number of potential links it could maximally have. For example, interaction types with higher specificity, such as parasitism, are expected to form networks with lower connectance than interaction types with lower specificity, such as seed dispersal. |
| Connectivity | The number and distribution of *links* in a *network* or *edges* in a *graph*. For example, if two networks have the same *size*, but one has fewer links, it has lower connectivity. Another example: if two networks have the same size and *degree*, but one is *nested* and the other is *modular*, they have different connectivity. |
| Criticality | Criticality point is the point where a small change in the average degree of a network (like adding or removing a link or node) causes a big shift in how the network behaves—like a system going from chaotic to stable. |
| Degree | A metric of *centrality*. The number of links that a *node* has in a *network*, or the number of *vertices* in a *graph*. For example, a bat species that visits several flower species in a network has higher degree than a bat species that visits few flower species. |
| Drawing | The visual representation of a *network* or *graph*. There are several methods for drawing networks and graphs, based on algorithms focused on highlighting different properties of the *system*. For instance, *bipartite* algorithms emphasize the different sets of nodes in the network, while energy-minimization algorithms emphasize the *centrality* of different *nodes* and the *modularity* of the network, and circular algorithms try to arrange the nodes in a circle or sphere. |
| Edge | Used in *graph* theory. A *relationship* between *vertices* of a graph. |
| Element | An entity that belongs to a *set* or a *system*. |
| Emergent property | A property that emerges from the assembly of a complex *network*. Emergent properties may be weak, when they exist by definition, or strong, when they emerge from the *system*. For example, if a bat-plant network is *nested*, *nestedness* is one of its weak emergent properties, while *robustness* to extinctions may be one of its strong emergent properties. |
| Foodweb | A *network* whose *links* represent trophic interactions (“who eats whom?”). For example, a network formed by bats and their predators, or bats and their prey. |
| Graph | Used in graph theory. A *set* of *vertices* (*elements*) and the *edges* (*relationships*) between those vertices. Graphs are abstract systems |

studied in pure mathematics. When a graph is studied in applied mathematics and represents a real-word system, it is called a *network*.

Layer — A set of *links* of one type in a *multilayer network*. For example, in a bat-plant network that contains two types of interactions, let us say frugivory and nectarivory, the frugivory links form one layer of the network.

Link — Used in *network* theory. A *relationship* between nodes of a network. For example, an interaction of pollination between a bat species and a plant species.

Modular — A *network* in which *modularity* predominates as a *topology*. For example, a pollination network in which different plant families form separate subgroups visited by different bat species.

Modularity — The *topology* of a *network* in terms of how many cohesive subgroups (i.e., *modules* or *communities*) it contains and how much *connectivity* there is between those modules.

Module — A subgroup of *nodes* in a *network* that are more densely (in a *binary* network) or strongly (in a *weighted* network) connected to one another than to other nodes of the same network. It may also refer to a subgroup of links in a network that share a very similar subgroup of vertices. For example, in a bat-plant network, a subgroup of bat species that feed on the same plant species.

Motif — A small, recurring link pattern within a network. It can be considered as a network substructure, with link arrangements that can convey social closure or bonding (such as triads), social centralization, and social-ecological motifs for a given shared resource, for instance, or social-ecological brokerage (when by engaging with multiple resources a social actor can share diverse ecological knowledge and resources to other actors).

Multilayer — A *network*, in which two *nodes* may be connected to one another by two or more types of *links*. For example, a network formed by bats and the plants that they visit to feed on fruits (link type 1) or nectar (link type 2).

Multipartite — A *network* that is divided into three or more *sets* of *nodes*, in which links may exist only between nodes of different sets. For example, a network formed by nectarivorous bats, the plant species visited by them, and the ectoparasites of those bats. Tripartite *foodwebs* are also called tritrophic networks.

Nested — A *network* in which *nestedness* predominates as a *topology*. For example, a pollination network, in which the plant species visited by a

| | |
|---|---|
| | bat species with lower *degree* are also visited by a bat species with higher degree. |
| Nestedness | The *topology* of a *network* in terms of how much the *links* of *nodes* with lower degree represent a subset of the links of nodes with higher degree. |
| Network | Used in *network* theory. A *set* of *nodes* (*elements*) and the *links* (*relationships*) between those nodes. For example, in Biology, real-word systems, such as ensembles formed by plants and pollinators, are modeled as networks. |
| Node | Used in *network* theory. An *element* of a *network*. For example, a bat species in a pollination network at the community level. |
| One-mode | See *unipartite*. |
| Path | A *set* of *links* in a *network* that connect two *nodes*. A *small path* is a geodesic, i.e., the smallest set of links between two nodes. |
| Projection | A *subnetwork* formed by one set of *nodes* from a *multipartite network*. For example, in a bat-plant network, if one builds a subnetwork with bat species only, in which two bat species are connected to one another by a *link* when they share at least one plant species, this is a *unipartite* projection. |
| Relationship | A *connection* between *elements* in a *system*, *network*, or *graph*. |
| Robust | A *network* that has the *emergent property* of *robustness*. |
| Robustness | An *emergent property* of some *networks* that suffer little change in *topology* when a given proportion of its *nodes* or *links* are removed. For example, a bat-plant network is robust, when it can lose several flower species without changing from *nestedness* into another topology. |
| Set | A group of *elements* in a *system*, *network*, or *graph*. |
| Size | The number of *nodes* in a *network*, or the number of *edges* in a *graph*. For example, the sum of bat and plant species in a pollination network. |
| Subnetwork | A subset of the *nodes* and *links* of a *network*. For example, in a bat-plant network that includes different plant families, if one builds a network with only the Piperaceae and the bat species that visit those plants, this is a *subnetwork*. |
| System | A *set* of *elements* and the *relationships* between those *elements*. *Networks* and *graphs* are kinds of systems. |

| | |
|---|---|
| Topology | The structure of a *network* or *graph* in terms of *degree*, *size*, and *connectivity*. For example, *nestedness* and *modularity* are two kinds of topology. |
| Two-mode | See *bipartite* network. |
| Unipartite | A *network* that contains only one *set* of *nodes*, in which *links* may exist between any nodes. For example, a network of social relationships between individuals within a bat population. Synonym: *one-mode network*. |
| Vertex | Used in *graph* theory. An *element* of a graph. |
| Weighted | A *link* to which some numerical weight is attributed. A *network* based not only on presence or absence of links, but also on the weight of those links, is called weighted. For example, the frequency of visits observed in the field of a given bat species to a given plant species may be used to attribute a weight to the link between them. |

**Table S4**. Most common SE nodes in network-bat studies (N>1).

| Nodes_EE | Count |
|---|---|
| Bats and plants | 24 |
| Bats and bat flies | 15 |
| Bat species and sampled areas | 5 |
| Bat species and bat flies | 4 |
| Bats and insects | 4 |
| Bats and roost | 3 |
| Mammals including bats | 3 |
| Bats and bat flies and fungi | 2 |
| Bats, Birds and Plants | 2 |
| Individuals of Desmodus rotundus | 2 |
| Individuals of Myotis lucifugus | 2 |
| Individuals of Myotis nattereri | 2 |
| Males and females of Rousettus aegyptiacus | 2 |

Supplementary Figures

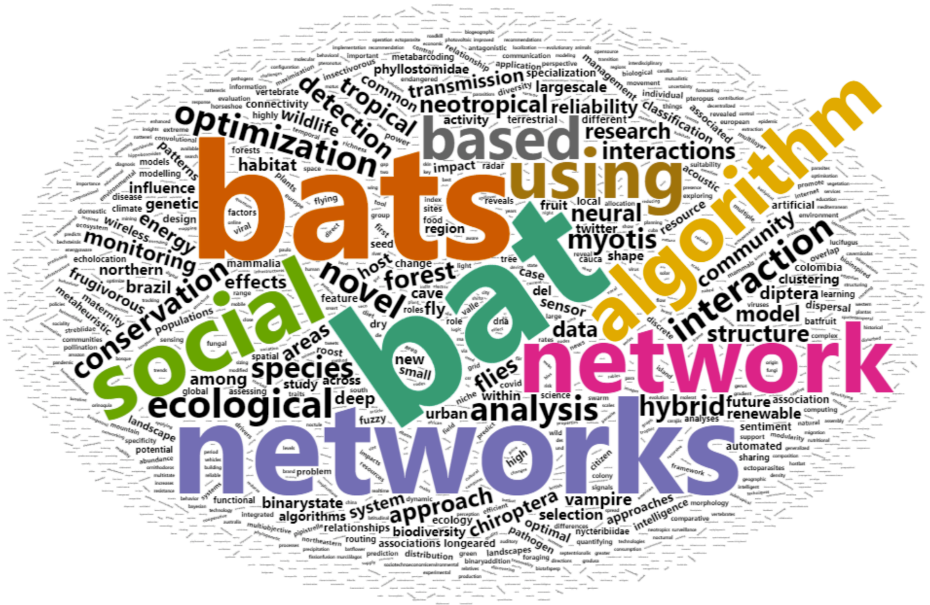

**Figure S1. Screening strategy -narrower:** socio* OR socia* OR ecolog* AND bat OR chiroptera AND network* OR graph* (322 Scopus) - Scopus_27_09_2024_with_social_keywords.csv. Code available as **pre-screening.R**

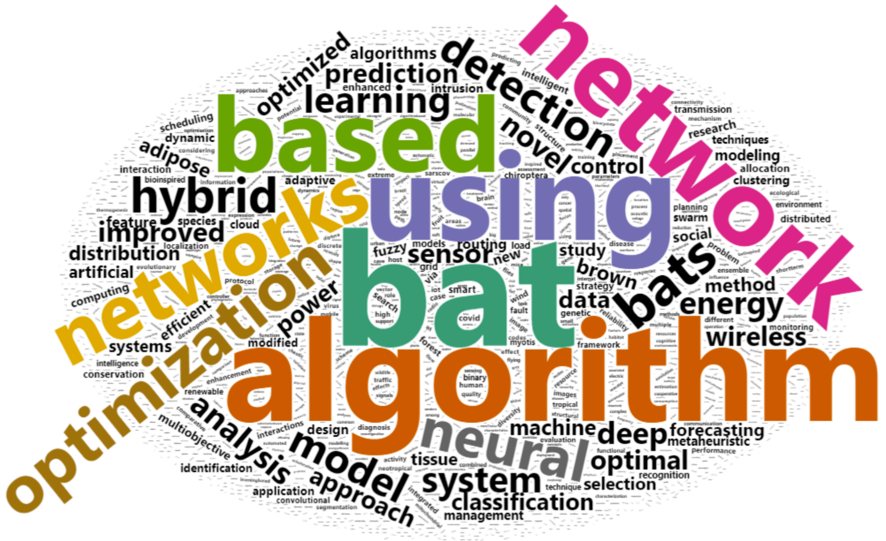

**Figure S2. Screening strategy -broader:** complementary comprehensive search using bat OR Chiroptera AND network* OR graph* (1,856 articles) - Scopus_26_09_2024.csv. Code available as **pre-screening.R**.

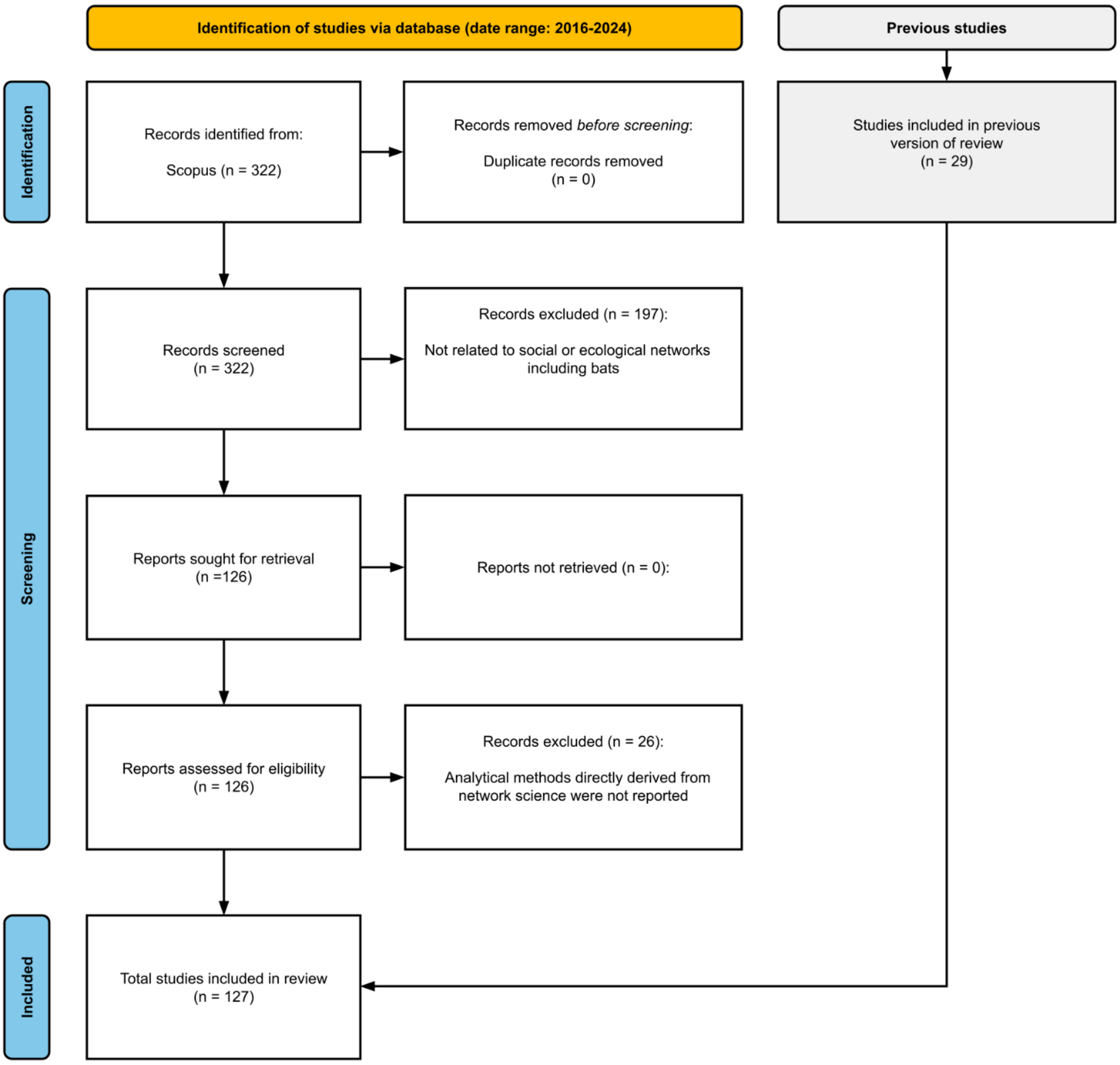

**Figure S3.** PRISMA chart, a visual representation of the stages of our bibliometric mapping used to review the literature around network studies focused on bats.

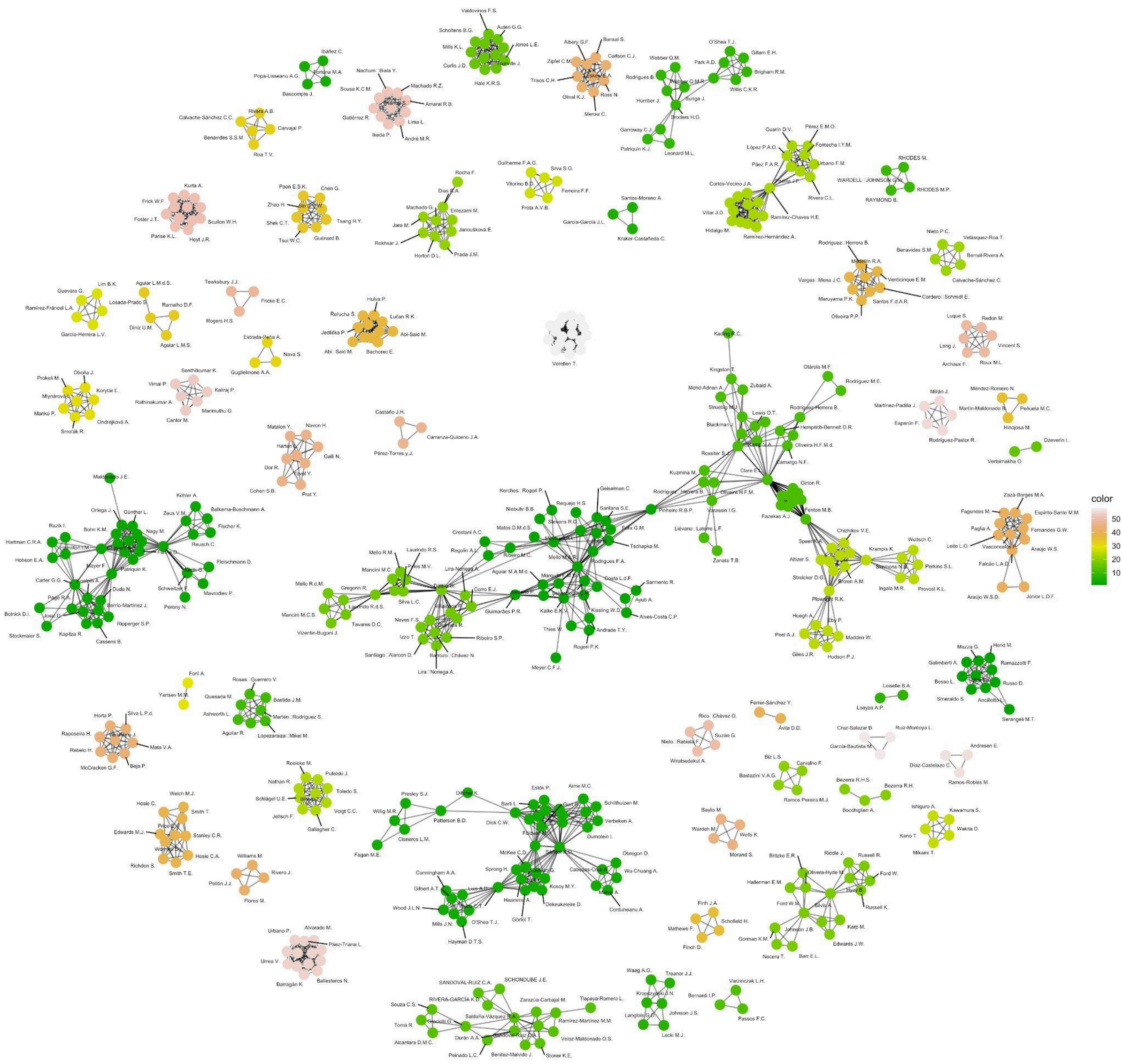

**Figure S4. Author (nodes) and their collaborations (links) revealed by the bibliographic mapping using a force-directed layout and label repelling to avoid figure noise**. Colour indicates the cluster membership through modularity analysis.

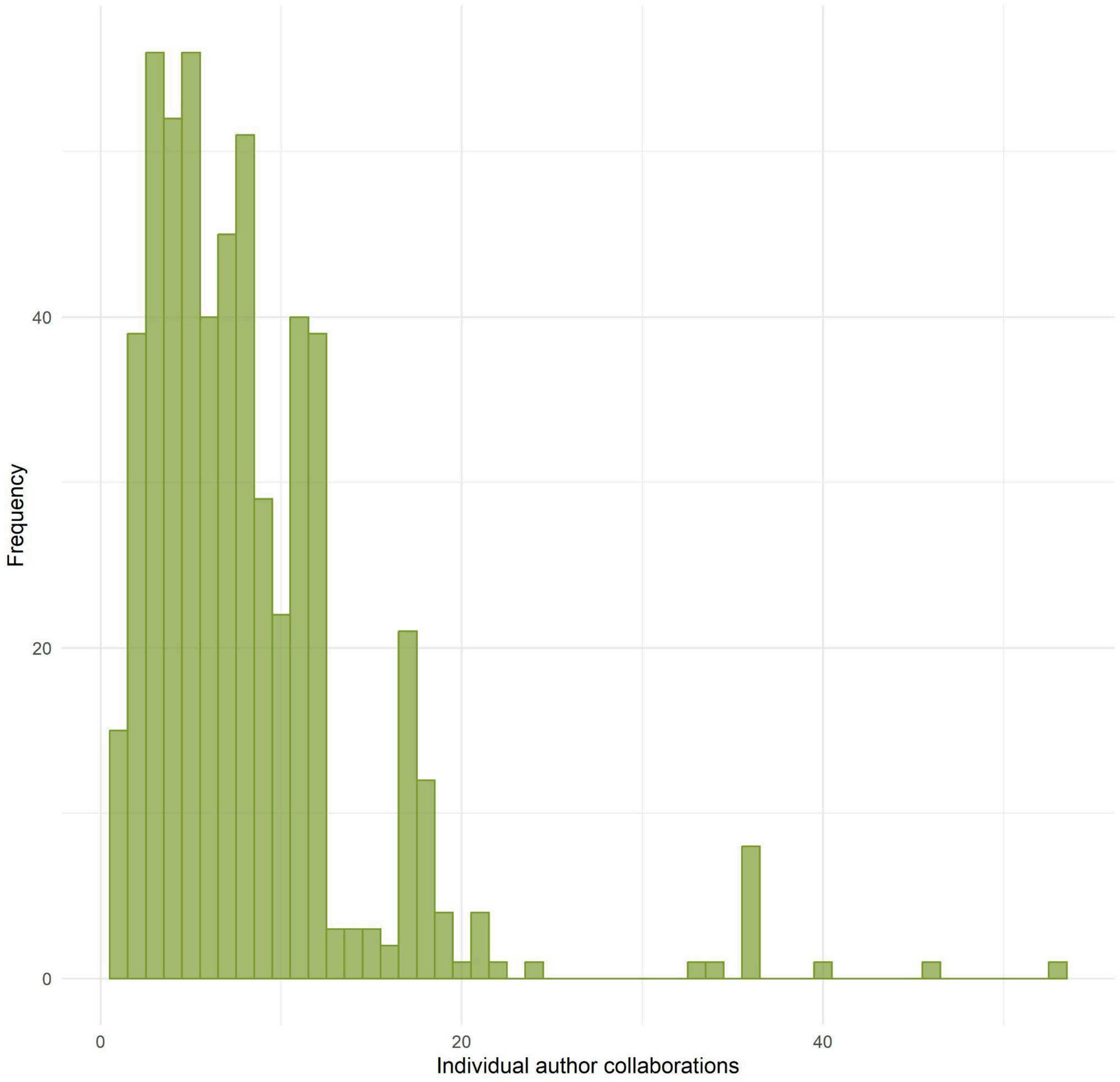

**Figure S5.** Author collaborations (degree distribution of the collaborator network) in bat studies with a network approach published between 2006-2024.

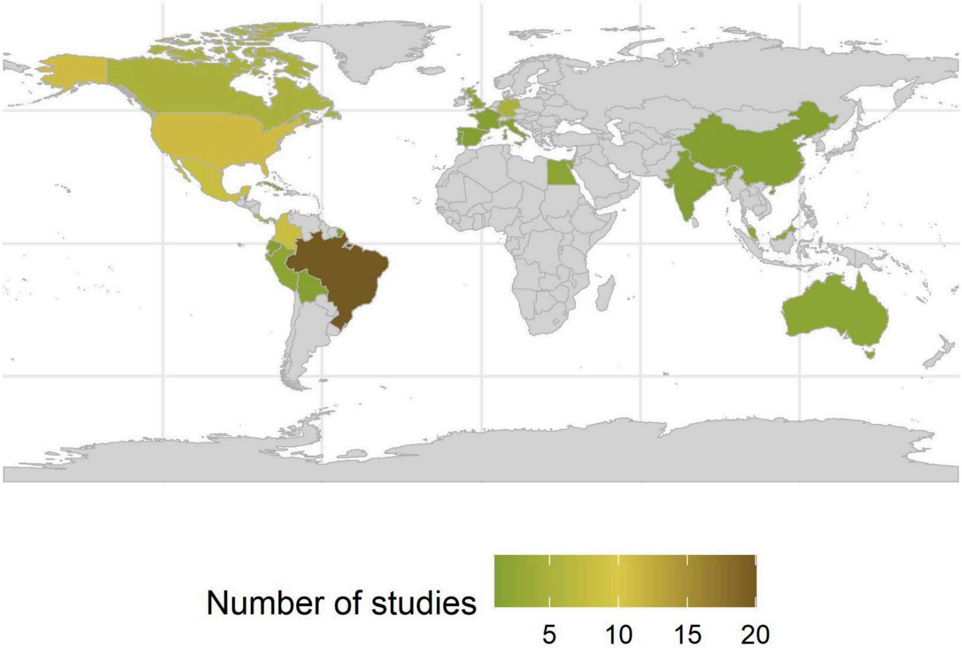

**Figure S6.** Geographical coverage of bat studies with a network approach found in our systematic review (2006-2024).
